# Optimized phenotype definitions boost GWAS power

**DOI:** 10.1101/2024.06.11.598562

**Authors:** Michael Zietz, Kathleen LaRow Brown, Undina Gisladottir, Nicholas P. Tatonetti

## Abstract

Complex diseases are among the central challenges facing the world, and genetics underlie a large fraction of the risk. Observational data, such as electronic health records (EHR), offer numerous advantages in the study of complex disease genetics. These include their large scale, cost-effectiveness, information on many different conditions, and future scalability with the widespread adoption of EHRs. Observational data, however, are challenging for research as they reflect various factors including the healthcare process and access to care, as well as broader societal effects like systemic biases. Here, we introduce MaxGCP, a novel phenotyping method designed to purify the genetic signal in observational data. Our approach optimizes a phenotype definition to maximize its coheritability with the complex trait of interest. We validated the method in simulations and applied it to real data analyses of stroke and Alzheimer’s disease. We found that MaxGCP improves genomewide association study (GWAS) power compared to conventional, single-code phenotype definitions. MaxGCP is a powerful tool for genetic discovery in observational data, and we anticipate that it will be broadly useful for studying complex diseases using observational data.

## 1 Introduction

The genetic basis of complex disease remains imperfectly characterized despite decades of theoretical and experimental advances and thousands of studies into the genetics of various complex diseases [1]. Piecing apart this causality will involve—among many other things—characterizing the associations between individual genetic variants and disease risk for millions of genetic variants and thousands of phenotypes. Observational data provide a practical source of information about complex diseases due to their abundance, representativeness, low cost to use, and the fact that they have already been collected. However, observational data bring additional challenges such as incompleteness, noise, and bias [2], which reduce study power [3].

By prioritizing genetic over environmental causes, careful phenotype definitions are one way to strengthen the genetic signal available in observational data [4, 5, 6]. Complex phenotypes are often correlated, leading researchers to analyze multiple phenotypes at once in the hopes of increasing study power [7]. However, combining related phenotypes in an optimal way is challenging, and previous work has dealt primarily with combining phenotypes that are known *a priori* to be related to the phenotype of interest [5, 6, 8]. Such approaches are inapplicable to phenome-wide analyses of biobank datasets, in which thousands of phenotypes are available. Moreover, by requiring features to be pre-selected, such methods are not able to incorporate weak genetic signals across large numbers of seemingly unrelated phenotypes. A method that can be applied to thousands of phenotypes without manual feature selection could increase power for phenome-wide studies.

Here we present MaxGCP, a new statistical method for defining phenotypes in GWAS. MaxGCP fits a linear combination of phenotypes to maximize coheritability with the phenotype of interest. As an example, a MaxGCP phenotype (*y*) for stroke could be defined as the linear combination of heart disease, hypertension, and other risk factors (e.g. *y* = 1.2 × heart disease − 0.1 × hypertension + …) that maximizes the coheritability between stroke and *y*. By boosting the genetic signal using information from other phenotypes, this method is able to reduce environmental noise and improve GWAS power.

## 2 Results

The goal of MaxGCP is to fit a phenotype *y* to maximize the coheritability between *y* and *z*, the target phenotype. In the previous example, *y* = 1.2×heart disease−0.1×hypertension+…, the features are heart disease, hypertension, etc., and the coefficients (i.e. 1.2, -0.1) are optimized by MaxGCP. A general overview of MaxGCP follows.

Suppose any phenotype, A, can be written as a sum of genetic and environmental components (i.e. *A* = *g*_*A*_ + *e*_*A*_). The coheritability between two phenotypes, A and B is

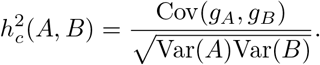

Define an index *y* as a linear combination of feature phenotypes *x*_1_, …, *x*_*m*_, (i.e. *y* = **x**^T^*β*). MaxGCP finds the 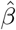 that maximizes coheritability between index *y* and target *z*. Because the index is linear, MaxGCP has an exact solution. Let **v** be a vector of genetic covariances between the features and the target (i.e. 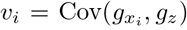). Let **P** be the covariance matrix of the features (i.e. *P*_*i*,*j*_ = Cov(*x*_*i*_, *x*_*j*_)). The optimal coefficient vector can be obtained by solving a linear system and is the following:

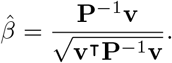

A derivation is provided in the methods.

### 2.1 Evaluation using simulated data

We first evaluated MaxGCP using simulated data. In simulation, effect sizes can be known exactly, making it easy to evaluate whether MaxGCP achieves its aims. We simulated 5000 realistic phenotypes (see Methods; Figure S1) and computed an optimal index for each phenotype using our method, MaxGCP. MaxGCP phenotypes were more heritable and coheritable than naively defined phenotypes (i.e. presence or absence of a single diagnosis code; Figure 1). This indicates that MaxGCP is able to prioritize specific genetic signals. When used in GWAS, MaxGCP phenotypes increased power compared to naive phenotypes (Figure 2, Table 1). MaxGCP’s performance depended on the quality of inputs used to fit the model. Excellent inputs (i.e., noiseless genetic covariance estimates) result in nearly optimal performance, while poor inputs (i.e., noisy estimates) result in worse performance.

**Table 1:** GWAS performance of each phenotype definition in simulation. Each reported metric is the median of per-phenotype values. A p-value cutoff of 0.05 was used to declare variants as predicted positives or negatives, and associations were compared to those correctly identified by the genetic component GWAS, making it perform perfectly by construction. Methods marked by an asterisk (*) are unobservable in real data.

| Method | Sensitivity | Specificity | Precision | F1 | AUROC |
| --- | --- | --- | --- | --- | --- |
| Genetic component* | 1.00 | 1.00 | 1.00 | 1.00 | 1.00 |
| Optimal linear* | 0.72 | 0.96 | 0.92 | 0.81 | 0.93 |
| MaxGCP (optimal) | 0.69 | 0.96 | 0.92 | 0.79 | 0.91 |
| MaxGCP (noisy) | 0.47 | 0.94 | 0.84 | 0.60 | 0.82 |
| Naive | 0.28 | 0.96 | 0.80 | 0.41 | 0.72 |
| Binary* | 0.37 | 0.96 | 0.85 | 0.52 | 0.79 |
| Liability* | 0.55 | 0.96 | 0.91 | 0.68 | 0.89 |

**Figure 1.**
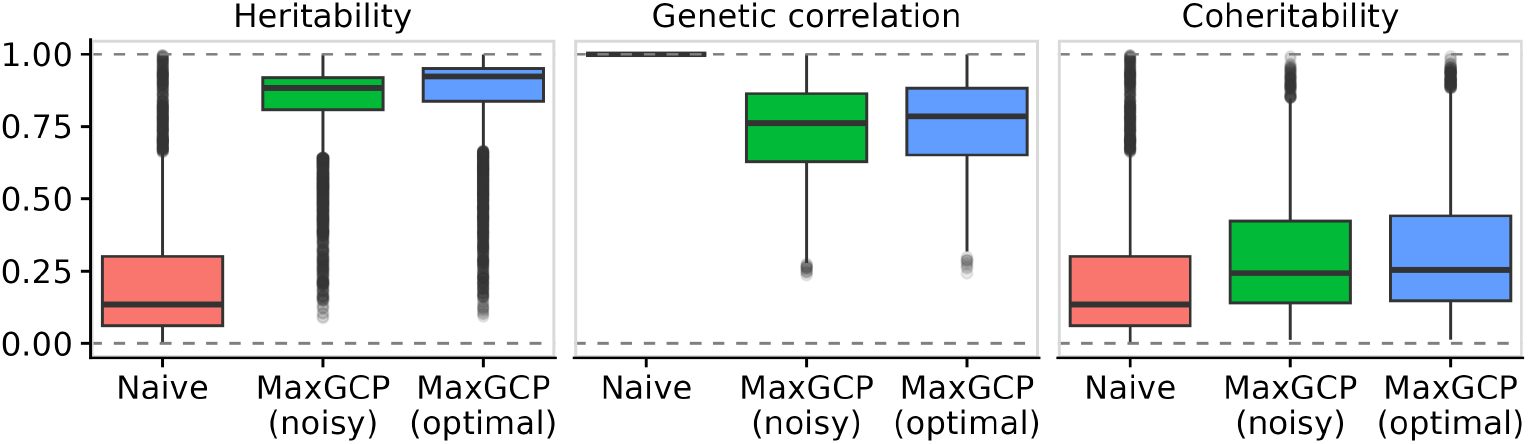
MaxGCP amplifies the genetic signal in simulated data. Shown is a comparison across phenotype definitions for 5000 simulated phenotypes. Naive definitions are single-code binary phenotypes with missingness. MaxGCP definitions are fit using either a noisy or noiseless (“optimal”) genetic covariance matrix. Genetic correlation and coheritability are computed relative to the naive definitions.

**Figure 2.**
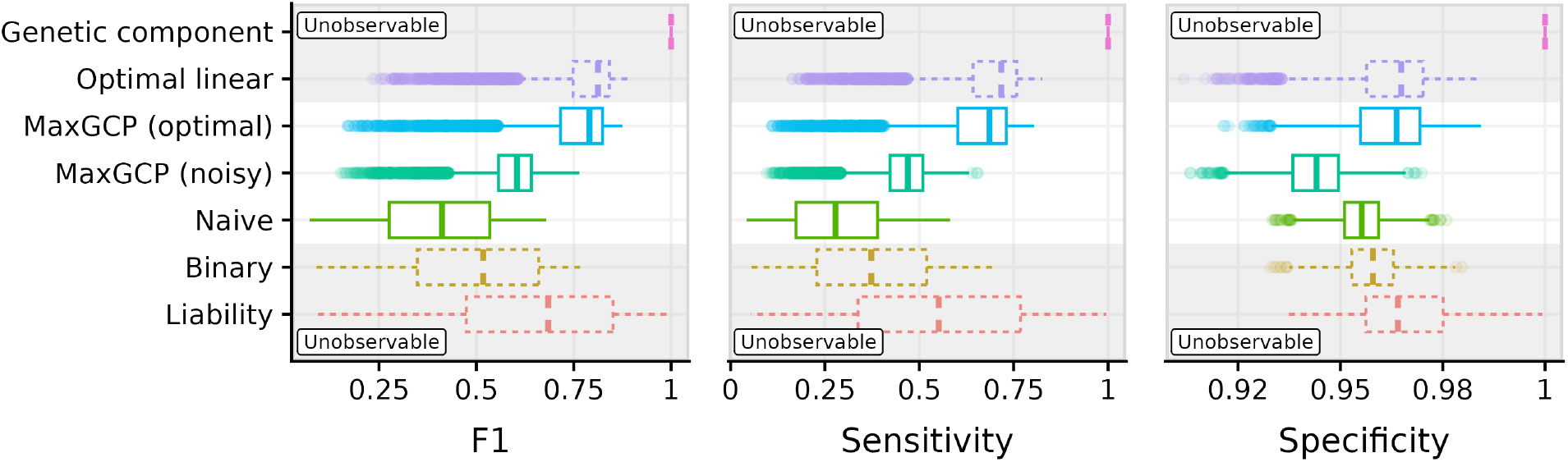
MaxGCP improves GWAS performance in simulated data. Shown are GWAS performance metrics for seven methods to define phenotypes in simulated data. Only naive and MaxGCP can be used in real data. These are indicated by the white strip through the middle of the plot. MaxGCP noisy and optimal provide bounds of reasonable performance, with the difference being the quality of genetic covariance estimates. Performance metrics are computed using a p-value cutoff of 0.05 to declare variants as positive or negative, and associations are compared to those correctly identified by the genetic component GWAS.

As a sensitivity analysis, we repeated this analysis using a variety of different simulation parameters (Figure 3). The results of this analysis are intuitive. Increasing the heritability of phenotypes increased GWAS power for all methods. Increasing the genetic correlation between phenotypes had a strong positive effect on MaxGCP GWAS but little effect on naive GWAS. Missingness (fraction of cases randomly switched to controls) reduced performance for all methods. Increasing variance in estimates of genetic covariance decreased MaxGCP performance and had no effect on naive phenotype GWASs, which do not take genetic covariance into account. Overall, the results of this sensitivity analysis were consistent with the previous results: (1) MaxGCP outperformed the naive phenotype (in sensitivity and F1, not in specificity), and (2) noise in the genetic covariance matrix degraded performance (for all metrics).

**Figure 3.**
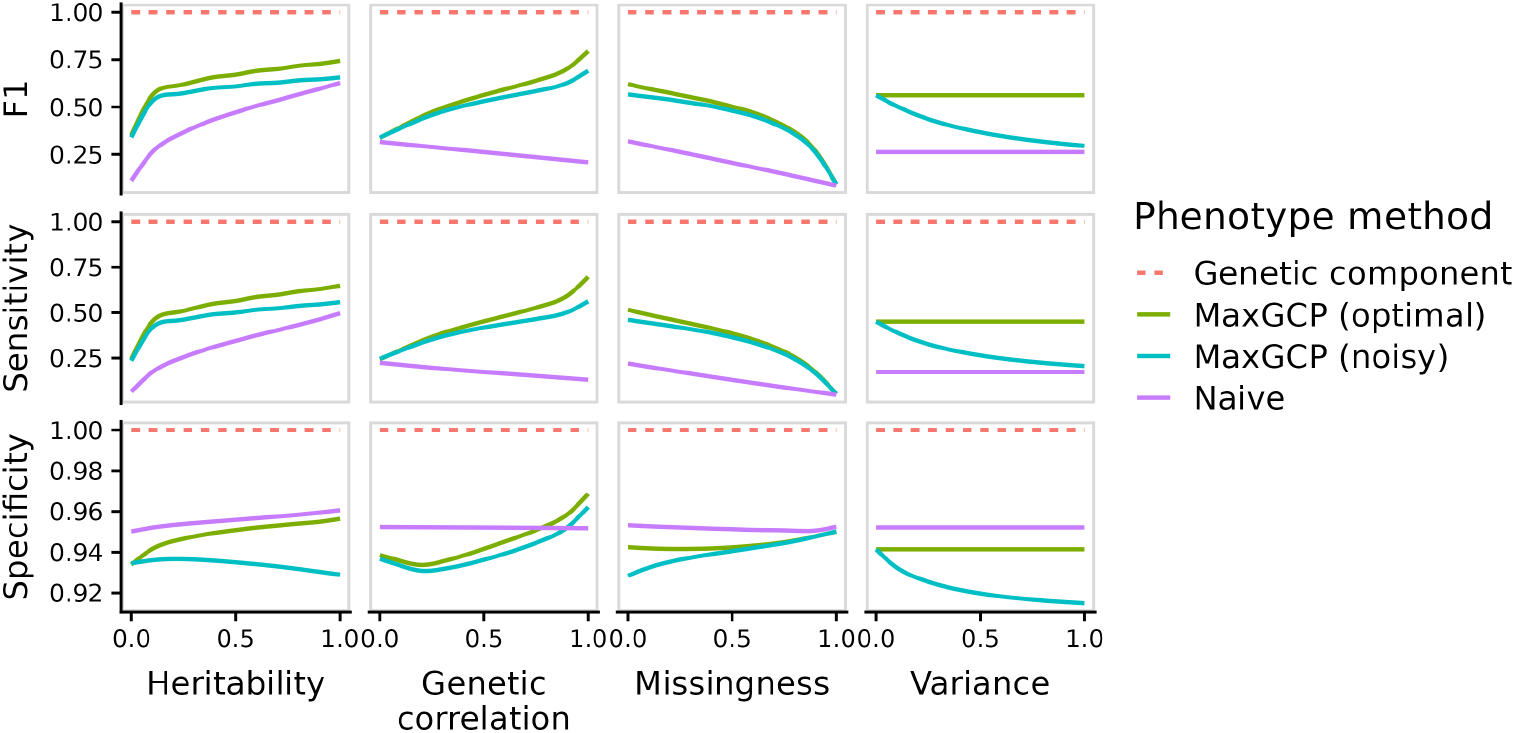
Sensitivity of GWAS performance to simulation parameters. Shown are marginal variations of parameters while all others are held constant at the realistic values. Each simulation parameter setting was simulated across 50 phenotypes and 100 replicates. GWAS performance was evaluated as before: relative to variants correctly identified by the genetic component GWAS.

### 2.2 Real data

Next, we evaluated MaxGCP using real phenotypes from the UK Biobank. This is a more challenging comparison because the true genetic architecture is not known. Our solution to this problem was to compare our GWAS results to external validation data, in the form of large, high-quality GWAS that do not include samples from the UK Biobank. We used two GWAS datasets for this task: MEGASTROKE (of which we used the “any stroke”, “ischemic stroke”, “cardioembolic stroke”, and “small vessel stroke” phenotypes) [9] and IGAP (Alzheimer’s disease) [10]. These studies used clinician-adjudicated phenotypes, which are more reliable than naive observational definitions (e.g. single instances of diagnosis codes from medical records) and, therefore, provide a high-quality reference for GWAS associations. As features for MaxGCP, we used the 50 most common ICD-10 codes in our cohort (see Methods). We ran GWAS using Plink 2 [11] and estimated genetic covariances using SumHer [12] (see Methods for further details).

MaxGCP improved sensitivity at the expense of specificity for all five phenotypes (Figure 4A, Table 2). However, there is an inherent trade-off between sensitivity and specificity, so we also compared at fixed thresholds. We evaluated sensitivity at 90% specificity and specificity at 10% sensitivity. MaxGCP performed well in this evaluation, with the full-cohort version performing best among all the methods for Alzheimer’s and three of the four stroke types (Figure 4B, Table 2).

**Table 2:** GWAS performance in real data. “P-value” results were computed by applying a p-value threshold of 0.05. “Fixed” results were computed by applying a fixed threshold to one metric (either 90% specificity or 10% sensitivity) and evaluating performance in the other. In both cases, predicted associations were compared to those made in the corresponding target GWAS (MEGASTROKE or IGAP). These values are the same as shown in Figure

| Target | Cohort | Phenotype | P-value |  | Fixed |  |
| --- | --- | --- | --- | --- | --- | --- |
|  |  |  | Sens. | Spec. | Sens. | Spec. |
| Any stroke | N=350k | MaxGCP | 0.120 | 0.910 | 0.130 | 0.929 |
|  |  | Naive | 0.065 | 0.947 | 0.114 | 0.913 |
|  | N=15k | MaxGCP | 0.055 | 0.947 | 0.101 | 0.901 |
|  |  | Naive | 0.057 | 0.948 | 0.105 | 0.906 |
| Ischemic stroke | N=350k | MaxGCP | 0.120 | 0.911 | 0.132 | 0.931 |
|  |  | Naive | 0.065 | 0.947 | 0.113 | 0.913 |
|  | N=15k | MaxGCP | 0.054 | 0.948 | 0.104 | 0.904 |
|  |  | Naive | 0.057 | 0.948 | 0.105 | 0.905 |
| Cardioembolic stroke | N=350k | MaxGCP | 0.075 | 0.931 | 0.108 | 0.909 |
|  |  | Naive | 0.059 | 0.947 | 0.107 | 0.906 |
|  | N=15k | MaxGCP | 0.052 | 0.947 | 0.098 | 0.898 |
|  |  | Naive | 0.055 | 0.948 | 0.102 | 0.902 |
| Small vessel stroke | N=350k | MaxGCP | 0.083 | 0.924 | 0.109 | 0.909 |
|  |  | Naive | 0.062 | 0.947 | 0.110 | 0.909 |
|  | N=15k | MaxGCP | 0.061 | 0.944 | 0.104 | 0.904 |
|  |  | Naive | 0.060 | 0.948 | 0.108 | 0.908 |
| Alzheimer's | N=350k | MaxGCP | 0.065 | 0.948 | 0.117 | 0.915 |
|  |  | Naive | 0.059 | 0.950 | 0.115 | 0.914 |
|  | N=15k | MaxGCP | 0.050 | 0.950 | 0.102 | 0.902 |
|  |  | Naive | 0.053 | 0.950 | 0.104 | 0.904 |

**Figure 4.**
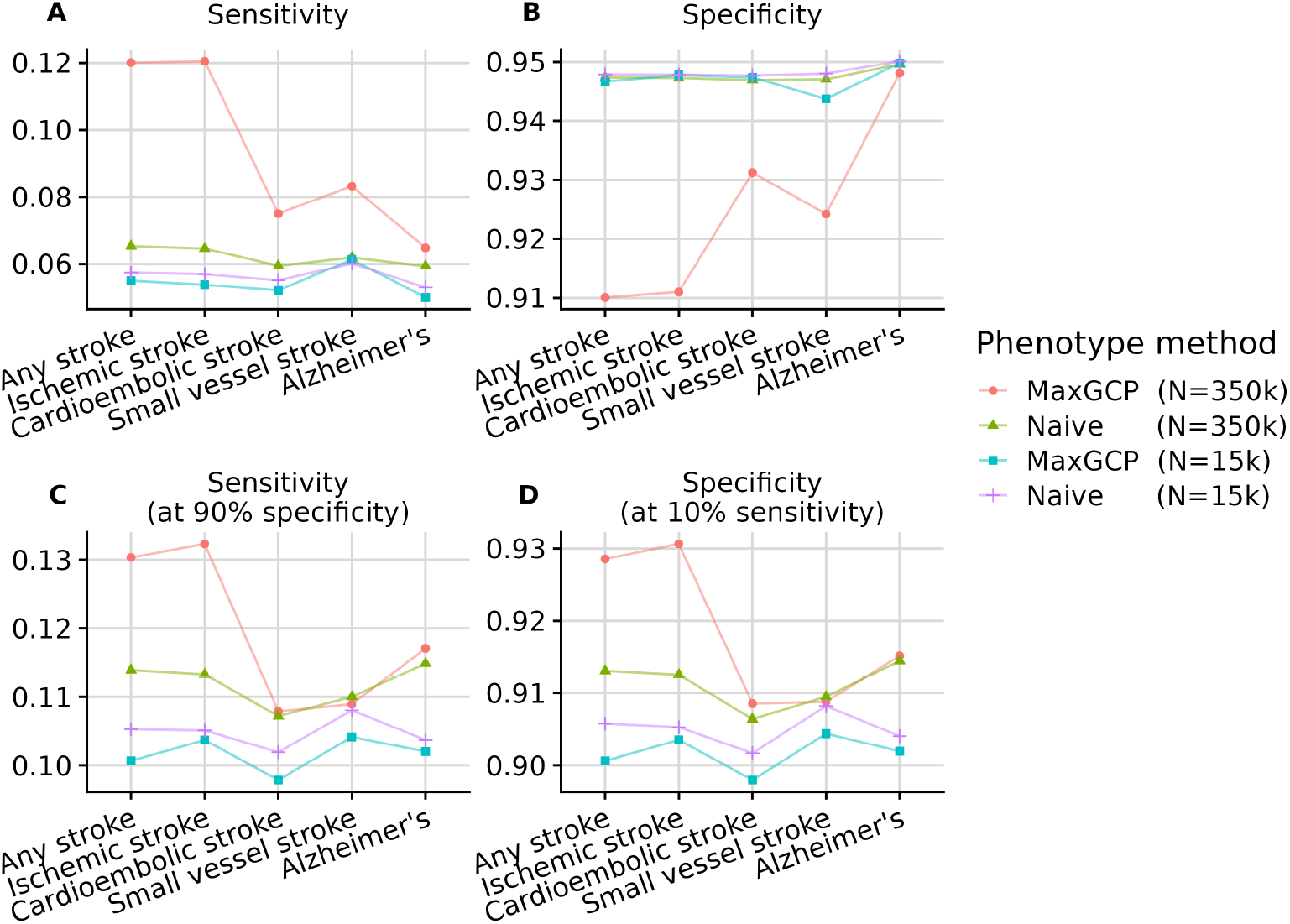
GWAS performance for real data. Phenotype definitions are compared in their ability to match variant associations with the MEGASTROKE and IGAP GWASs. Each analysis was performed in both the full and subsampled cohorts (N=350k and N=15k, respectively). **A** Sensitivity and specificity are evaluated using a single p-value threshold (*p <* 0.05). **B** Sensitivity and specificity are evaluated at 90% specificity and 10% sensitivity, respectively.

Overall, we draw the same conclusions as in the simulation. First, MaxGCP can improve GWAS sensitivity. Second, the quality of input data affects MaxGCP’s performance. MaxGCP showed excellent performance in the full cohort, but it underperformed in the subsampled cohort, where estimates of genetic covariance are less accurate. Third, MaxGCP improves sensitivity at the expense of specificity. Fourth, MaxGCP improves sensitivity more than it lowers specificity, meaning overall performance is generally superior (e.g. when comparing methods at equal specificity).

MaxGCP prioritizes relevant features, including known risk factors like heart disease for stroke and type 2 diabetes for Alzheimer’s (Figure 5 and Supplementary Table 3). The top features for stroke phenotypes were cerebral infarction itself (ICD-10 code I63), acute ischemic heart disease (code I24), and pulmonary heart diseases (I27). For Alzheimer’s disease, the top MaxGCP features were Alzheimer’s disease (G30), pulmonary heart diseases (I27), and type 1 diabetes (E10), with the latter two having negative coefficients.

**Figure 5.**
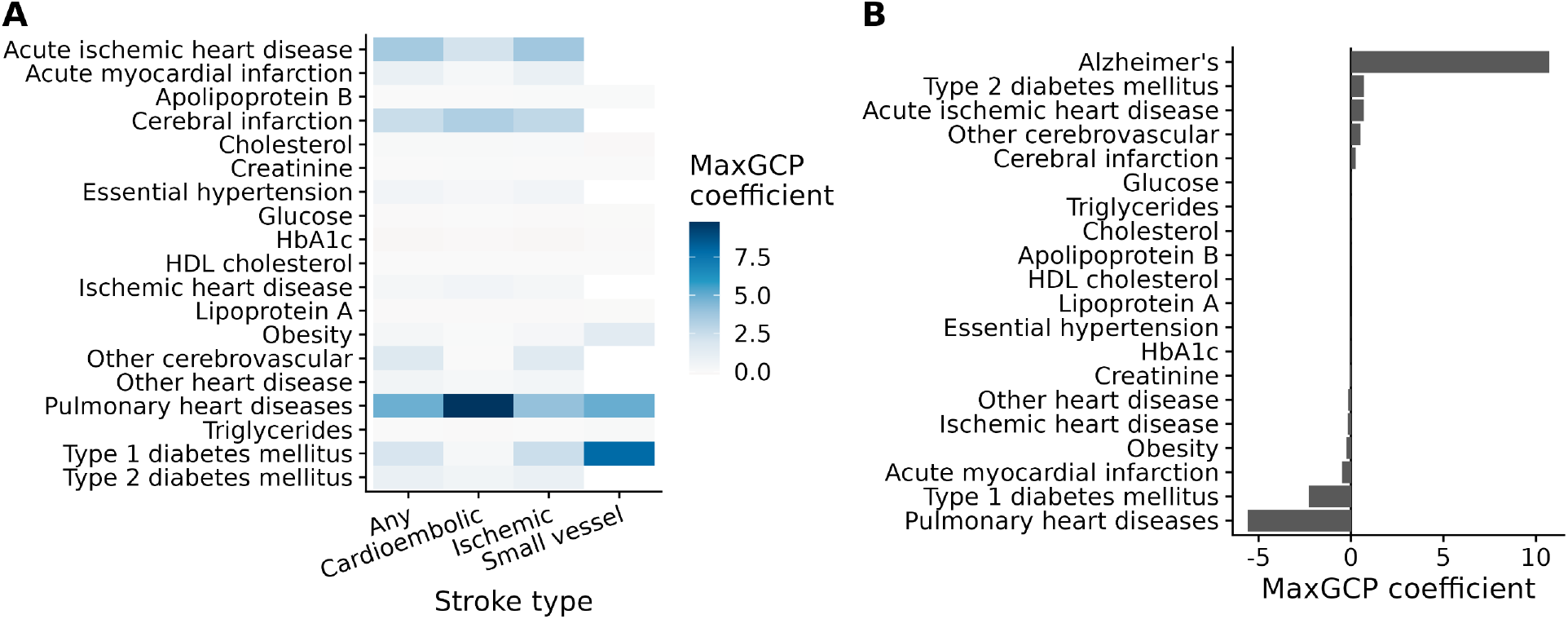
Top MaxGCP coefficients in real data. **A**. Top coefficients for MEGASTROKE phenotypes. **B**. Top coefficients for Alzheimer’s disease. Shown are the fitted coefficient values of the MaxGCP model. The corresponding MaxGCP phenotypes are linear combinations of the features with the shown coefficients. Most features received essentially zero weight, so we selectively visualize top coefficients along with known risk factors to present an interpretable view into MaxGCP.

**Figure 6.**
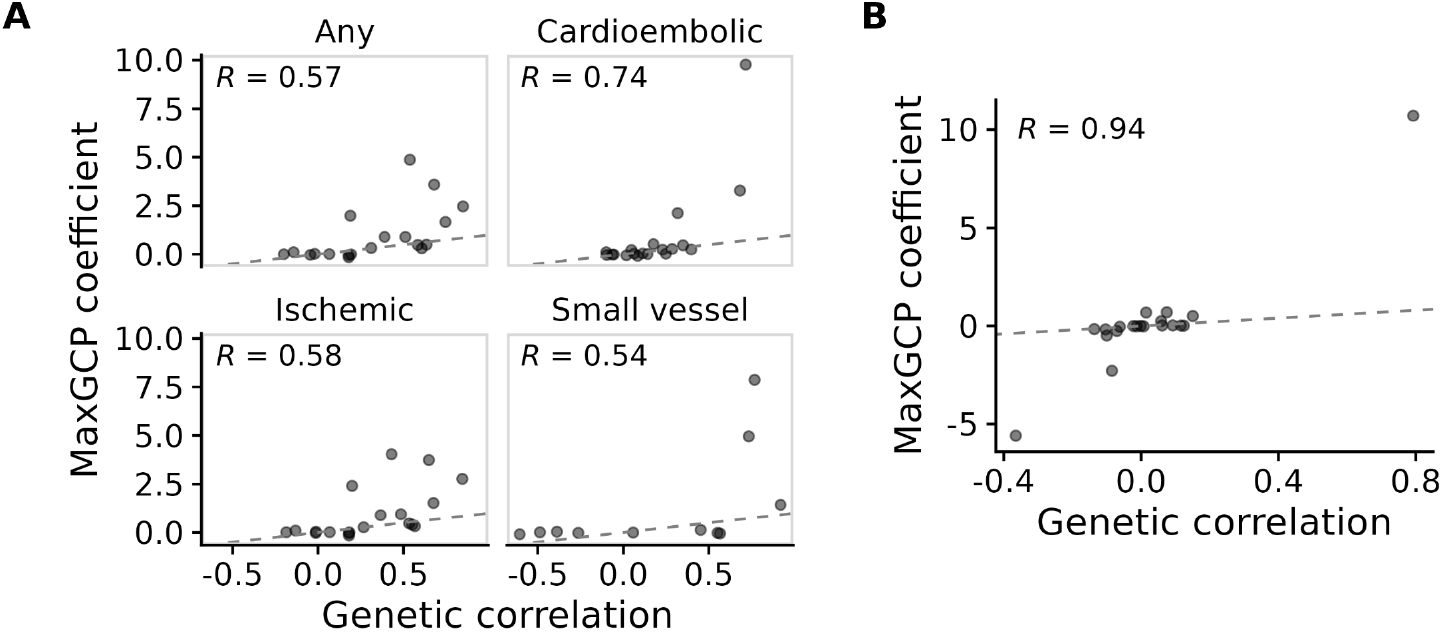
Genetic covariance vs MaxGCP coefficients in real data. Comparison of each feature phenotype’s genetic correlation with the target to the coefficient it received in the MaxGCP phenotype. **A**. MEGASTROKE phenotypes. **B**. Alzheimer’s disease. Each point represents a single feature phenotype. The y-axis represents the coefficient it received in the MaxGCP phenotype, while the x-axis represents the estimated genetic correlation between the target phenotype (i.e. stroke on the left, Alzheimer’s on the right) and the feature phenotype. Shown are the same features as in Figure 5 and Table 3.

MaxGCP coefficients are driven by both genetic and phenotypic covariance. Genetic covariance estimates were strongly, though imperfectly, correlated to MaxGCP coefficients (Pearson correlations between 0.54 and 0.94; Figure 6). In other words, the MaxGCP coefficient for a feature (e.g. hypertension) is predicted by the strength of genetic covariance between the feature and the target (e.g. Alzheimer’s disease). The fact that this correlation is not perfect suggests that MaxGCP correctly includes information about phenotypic covariances, which in effect minimizes phenotypic variance while maximizing genetic covariance.

MaxGCP GWAS identified target peaks that are not present in the naive GWAS but are in the target (Figures S2 and S3). If MaxGCP functions as expected, then a Manhattan plot of its GWAS results should look more similar to the target (e.g. MEGASTROKE) than the naive phenotype itself (e.g. ICD-10: I63). Moreover, MaxGCP results should not resemble the results for a top feature more than they resemble the target, as this would suggest that MaxGCP increases the number of hits by including false positives. Comparisons of Manhattan plots from this analysis confirm that MaxGCP is performing correctly, providing additional evidence that MaxGCP correctly identifies phenotype-specific information.

### 2.3 Comparison to MTAG

We compared MaxGCP’s performance in real data to MTAG. We were not able to perform a full comparison for two reasons. First, MTAG requires pairwise LD score regressions among all feature phenotypes. As a consequence, the number of LD score regressions required for MTAG grows quadratically in the number of features used. MaxGCP, on the other hand, requires only a linear number. Because of this, the MTAG comparison was not computationally feasible for us using large numbers of features (e.g. ≥ 50). Second, because the IGAP Alzheimer’s disease GWAS summary statistics do not include allele frequencies, we excluded Alzheimer’s from this comparison.

For this comparison, we used four different phenotype methods: the naive phenotype (a single ICD code), MTAG (with 10 features), MaxGCP (with 10 features), and MaxGCP (unrestricted features). The unrestricted MaxGCP was the model used previously, where features are selected for each phenotype using quality control only. For the fixed feature comparisons, we selected the 10 features in decreasing order of their absolute genetic correlation to the target phenotype. Because the p-value cutoff plays such a large role in performance in this comparison, we evaluated across a range of p-value cutoffs.

We found that MaxGCP performs similarly to or better than the naive phenotype throughout this comparison. Without restricting the number of features used, MaxGCP outperformed MTAG throughout this comparison. With only 10 features, MaxGCP performed similarly or slightly better than MTAG in all comparisons. For example, with a p-value cutoff of 0.05, MTAG outperformed MaxGCP-10 for the All stroke phenotype (MTAG sensitivity at 95% specificity was 0.089, MaxGCP was 0.085) when using 10 features for both methods. However, with a p-value cutoff of 0.005, MaxGCP-10 performed better in the same metric (MTAG: 0.137, MaxGCP 0.144).

Figure 7 shows the results of this comparison. Overall, MaxGCP was the best performing method, particularly when unrestricted in the features it could use. MTAG performed well for all stroke and acute ischemic stroke, but underperformed for cardioembolic stroke. Compared to MTAG, MaxGCP has the advantages of significantly better computational complexity and no dependence on allele frequency information. This allowed us to use far more features for MaxGCP than for MTAG, leading to improved performance and reduced computation time.

**Figure 7.**
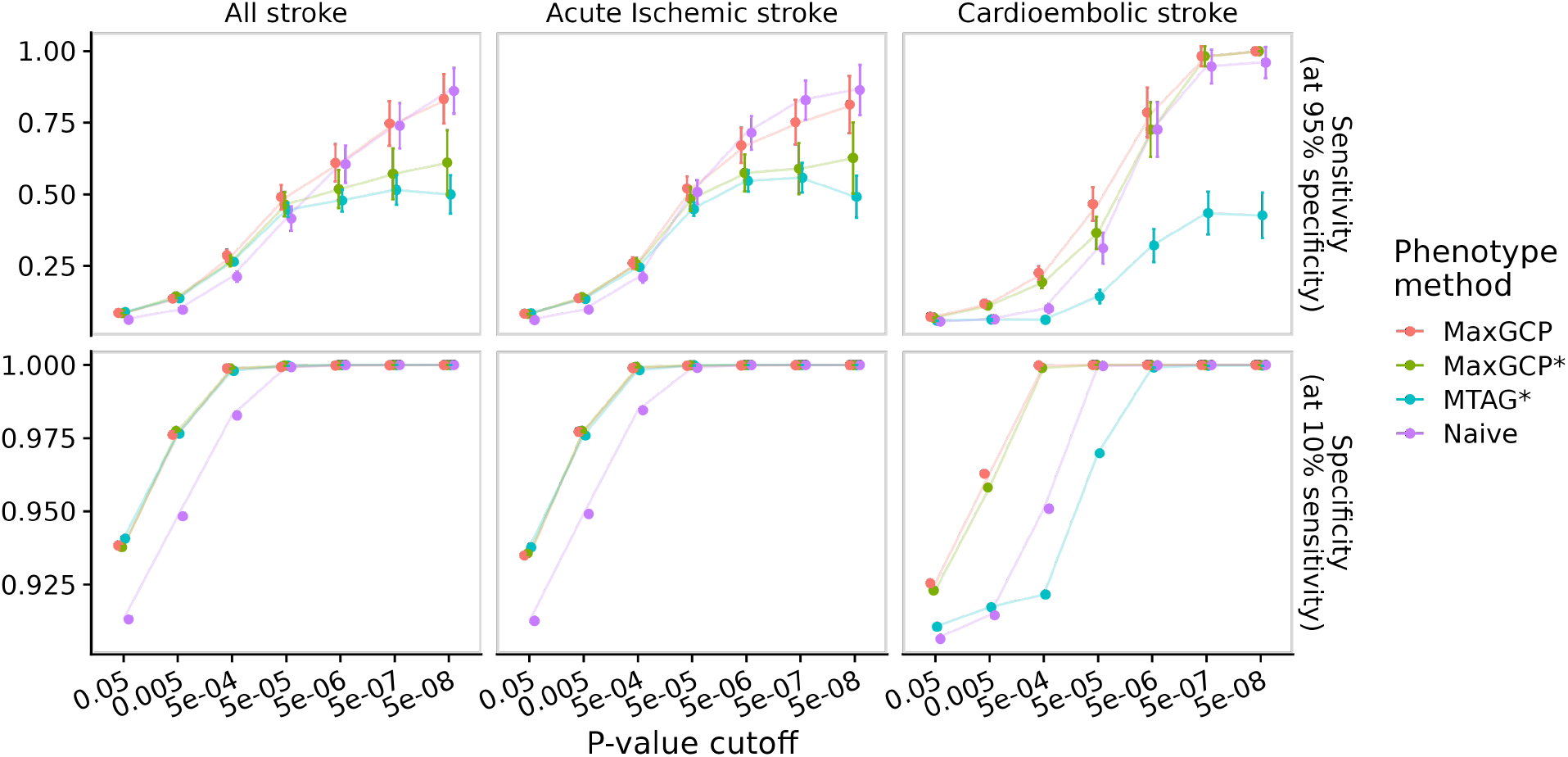
Comparison between MTAG and MaxGCP. Shown are the sensitivities and specificities of MaxGCP and MTAG in three phenotype GWAS. Sensitivity and specificity are evaluated at 90% specificity and 10% sensitivity, respectively, to ensure a fair comparison. Four phenotype methods are compared: MaxGCP (unrestricted), MaxGCP* (using 10 features), MTAG* (using 10 features), and Naive (a single ICD-10 code).

## 3 Discussion

Here we described MaxGCP, a method for defining phenotypes in observational data. MaxGCP takes two inputs—a phenotypic covariance matrix (feature phenotypes × feature phenotypes) and a vector of genetic covariance estimates (feature phenotypes × the target phenotype)—and it fits a linear combination to maximize coheritability with a target phenotype. Our analysis used both real and simulated data and provided two takeaways. First, MaxGCP improves the performance of GWAS. It increased power throughout our comparisons. While it reduces specificity, it increases sensitivity more than it reduces specificity, meaning that, overall, it performs better. Second, MaxGCP’s performance depends on the quality of inputs used. Given excellent genetic covariance estimates, MaxGCP can nearly double sensitivity (Figure 4A). However, MaxGCP can reduce power given poor estimates, meaning that quality control is critical when applying MaxGCP.

Phenotype-specificity is the key feature that distinguishes MaxGCP from most previous methods. Large, multi-phenotype datasets like the UK Biobank include many phenotypes that are not obviously related to one another (e.g. motor vehicle collisions and hair color). Previous methods like MaxH also optimize a linear phenotype definition for genetic signal, but they do not restrict to a specific genetic signal. A nonspecific method like MaxH cannot be naively applied to data like the UK Biobank, because a maximally heritable phenotype definition may not have any relation to the specific phenotype under consideration. For example, it may be the case that a combination of eye color and height produces the most heritable phenotype definition in a dataset, but this is not helpful for a researcher interested in the genetics of stroke. Unlike previous methods such as MaxH, MaxGCP can be applied without pre-selecting features based on relevance.

MaxGCP can be applied broadly, but it’s application merits some consideration. First, the quality of inputs is very important to MaxGCP performance. In particular, this means that only high-quality genetic covariance estimates should be used. Estimates that imply out-of-bounds values (e.g. heritability less than zero) or have large standard errors (e.g. *>* 2), should be excluded. In our analysis of real data, we used the most common ICD codes and large cohorts to give reliable genetic covariance estimates, and we excluded features according to the above criteria. However, applying MaxGCP to thousands of features without additional or more stringent quality control may lead to poor fits due to the combined effects of many noisy estimates. There are likely additional filters, adjustments, and quality control steps that could be used to further improve performance in such cases. Overall, researchers applying MaxGCP should pay careful attention to the quality of genetic covariance estimates used as inputs to the model and should remove features that do not have high-quality estimates.

We compared MaxGCP to MTAG, a previous method that is phenotype-specific. This comparison was not straight-forward due to differences in the methods. For one, the MTAG software does not separate features and targets in the same way MaxGCP does. When running MTAG, one specifies a set of GWAS summary statistic files, and MTAG generates MTAG summary statistics corresponding to every input file. This means that MTAG must compute pairwise LD score regressions among all phenotypes. Conversely, MaxGCP only requires genetic correlation estimates between the features and target, so the number of LD score regressions needed grows linearly, not quadratically, in the number of features.

Due to high computational costs, MTAG could not be run for large numbers (≥ 50) of phenotypes in this study. This particularly complicated the analysis because we needed to select which features should be included. For example, when the target phenotype was acute ischemic stroke, the features were the 10 phenotypes with the largest absolute genetic correlation to the target (i.e. acute ischemic stroke). While MaxGCP coefficients are correlated with genetic covariance estimates, this is not necessarily an optimal feature selection mechanism. Overall, we found that MaxGCP performed similarly to or better than MTAG. In addition, MaxGCP is faster, with its main computational cost growing linearly in the number of features, while MTAG grows quadratically due to pairwise LD score regressions.

MaxGCP relies on estimated genetic covariances, which may be biased by indirect genetic effects. For example, consider the relationship between type 2 diabetes and the ZIP code that one inhabits. ZIP code does not have a genetic basis, but neighborhood segregation along ancestry lines creates an association between genetics and ZIP code. As a consequence, a study which does not fully adjust for the effects of ancestry could find a genetic correlation between ZIP code and type 2 diabetes, even though the relationship is not causally genetic. MaxGCP coefficients are correlated with genetic covariance estimates, so biases like this will bias MaxGCP. Future work could address this directly. To give an example, a method could be devised in which MaxGCP penalizes genetic covariance estimates that are likely biased by population stratification (e.g. approximated via LD score regression intercept).

As is a common approach in GWAS, we primarily considered binarized variant associations using p-value cutoffs. Here, we were interested in validating the method, so our choice of p-value cutoff was arbitrary. We used a larger (i.e. more lenient) p-value cutoff (0.05) to ensure that enough hits were present to evaluate the method. Accordingly, the sensitivity and specificity values reported here may not reflect performance under typical GWAS cutoffs (i.e. 5 × 10^−8^). However, as shown in various studies over the last 15 years, the majority of SNP heritability is not captured by genome-wide significant variants but by many variants with small effects that do not reach statistical significance in current cohorts [13, 14]. Thus, while we use more lenient p-value cutoffs, our results may still be informative about the variants/effects underlying the true genetic architecture of the phenotypes.

A major limitation of the current analysis is that the reference GWAS are not well-captured by either naive or MaxGCP phenotypes (i.e. the overall sensitivity was low). We directly compared associated variants, but this is far from a perfect method of evaluation. In particular, differences in sequencing technology and imputation can lead to differences in GWAS results. As a potential future direction, we could consider whether GWAS associations are colocated with one another, as this would provide a more robust evaluation of the method and reduce the effects of unrelated technical choices in the production of reference GWAS. In future work, we also plan to evaluate MaxGCP using a more diverse set of phenotypes, as the current evaluation was limited to stroke and Alzheimer’s disease.

## 4 Methods

MaxGCP seeks to boost the power of GWAS by prioritizing the genetic component of a phenotype definition. To simplify this work, we focus on linear combinations of feature phenotypes, which we refer to here as indices. The heritability, coheritability, and genetic correlation of an index can be written in terms of the feature phenotypes. This fact allows us to optimize coheritability efficiently. Define an index (*y*) as a linear combination of (feature) phenotypes (x_1_, …, x_*m*_), where *β* is an arbitrary coefficient vector.

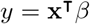

Denote the genetic and phenotypic covariance matrices of the feature phenotypes as **G** and **P**, respectively (i.e. 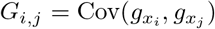 and *P*_*i*,*j*_ = Cov(*x*_*i*_, *x*_*j*_)). Denote the vector of genetic covariances between *z* and each feature phenotype as **v** (i.e. 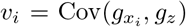). Using this notation, the definitions of heritability, genetic correlation, and coheritability are as follows:

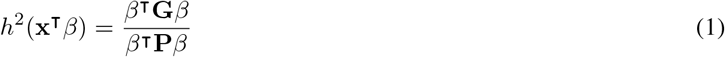

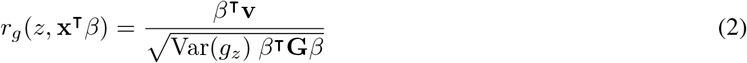

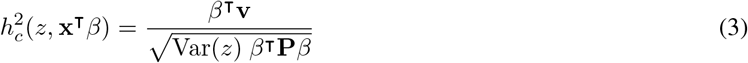

### 4.1 Coheritability optimization

To boost the genetic signal of an index, we seek to amplify genetic covariance while minimizing phenotypic variance. Coheritability is the appropriate objective for this task, as it is proportional to genetic covariance and inversely proportional to phenotypic variance. We seek to find coefficients that maximize the coheritability between *y* and *z* (i.e. 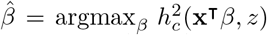). Note that the coheritability does not change based on a scaling of *β*. Accordingly, we can require that *β* be scaled so that the resulting index has unit variance. This allows us to simplify the problem to the following: maximize (w.r.t. *β*) *β*^T^**v** subject to *β*^T^**P***β* = 1. This can be solved using a Lagrange multiplier approach.

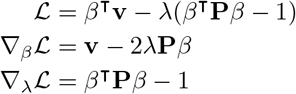

We set the gradient equal to zero and solve.

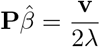

Applying the constraint, we can find *λ*:

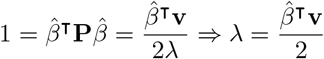

This allows us to rewrite the gradient equation:

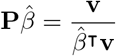

The matrix **P** is positive definite and can be inverted. We scale the result to ensure the index has unit variance, and obtain the solution as follows:

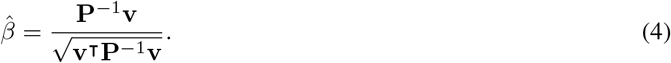

To summarize, a set of coefficients 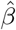 that maximize the coheritability between **x**^T^*β* and an arbitrary phenotype *z* can be determined by solving the linear system **P***β* = **v** and scaling the resulting coefficients by 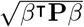.

### 4.2 Implementation

As shown above, MaxGCP is a straightforward operation that can be computed by libraries in nearly every programming language. We implemented this in a Python package [15] in order to pair the method with related functionality. Specifically, we added summarization functions that compute the heritability, genetic correlation, coheritability, and genetic loss given fitted coefficients, using Equations 1, 2, and 3, as well as methods to optimize heritability or genetic correlation instead of coheritability.

### 4.3 Simulation analysis

Real genotypes from the UK Biobank—10,000 (White British, QC-passing) individuals and 100,000 (imputed, HapMap3, QC-passing) variants—were used as the foundation for the simulation. We simulated 100 sets of 50 phenotypes, each with 5000 cases and controls, with a 1% mean heritability, 50% of variants having shared effects, zero mean phenotypic covariance, and 10% missingness using PhenotypeSimulator [16]. These parameters were chosen to simulate realistic phenotypes (Figure S1). For comparisons, positives were defined as variants simulated to be causal that were associated with nominal statistical significance (*p <* 0.05) in the genetic component GWAS. Negatives were defined as the opposite: non-causal variants with no statistically significant association detected in the genetic component GWAS. We qualified on the genetic component GWAS because this is the optimal GWAS phenotype and allows us to isolate the relevant performance of different methods from undesired randomness in the simulation.

We compared seven phenotype definitions in the simulation. The first two represent the simulation—the simulated continuous phenotypic liability (“liability”), and the binarized liability (“binary”)—and are not observable in real data. The binarized liability with missingness (“naive”) represents our best attempt at a realistic phenotype. MaxGCP models were fit using both noisy and noiseless genetic covariance matrices (“MaxGCP noisy” and “MaxGCP optimal”). Finally, we included two phenotypes that can only be observed in simulation. The optimal linear combination of phenotypes (“optimal linear”) was determined by directly regressing genetic components against phenotypes, and it represents the upper bound for the performance of a linear phenotype definition. The genetic component of each phenotype (“genetic component”) is included as it defines perfect performance.

### 4.4 Real data analysis

We used real data from the UK Biobank. Because we used the data to estimate genetic covariances, we applied the following QC filters: only White British individuals, excluded mismatched sex, outliers for heterozygosity or missingness, sex chromosome aneuploidy, relatedness to other participants, or those not included in the computation of genetic principal components. We used only imputed HapMap3 variants and excluded variants with high missingness. For feature phenotypes, we used ICD-10 diagnoses. We used SumHer [12] to estimate the genetic covariance between phenotypes. The GWAS inputs to SumHer were computed using Plink 2 with age, sex, and 10 genetic principal components as covariates. For each MaxGCP phenotype, we included only the 50 most common ICD-10 codes in our cohort as features. After estimating genetic covariances between each feature and each target, we excluded features that had invalid estimates for a given target. Extensive details about data collection and preparation are available in the supplement.

## 5 Supplementary Materials

### 5.1 Processing of UK Biobank data

To begin, we selected a cohort using only White British individuals to reduce the effects of population structure on our analysis. Next, we removed individuals whose data were flagged for various reasons as being potentially flawed or erroneous. Specifically, we removed individuals whose genetic sex was mismatched with their self-reported sex, individuals who were outliers for heterozygosity or missingness (defined by the UK Biobank using the outlier detection algorithm, *abberant* [17]), individuals with ten or more third-degree relatives in the UK Biobank, individuals with sex chromosome aneuploidy, and we restricted to individuals used in the computation of genetic principal components by the UK Biobank. Finally, we restricted to individuals with diagnosis data available, as described below.

We gathered phenotypic data for this cohort using the International Statistical Classification of Diseases and Related Health Problems, 10th revision (ICD-10) codes. In the UK Biobank, these can be obtained from six different data fields, and we included them all. These fields are hospital inpatient ICD-9 codes, hospital inpatient ICD-10 codes, self-reported non-cancer illness codes, primary cause of death, secondary cause of death, and general practitioner outpatient diagnoses. We used mappings provided by the UK Biobank to convert each coding to ICD-10. Only codes with at least 100 observations were retained. Applying the above QC and filtering procedure resulted in 1238 binary phenotypes and 342,350 samples. To save computation time, we restricted our analysis to HapMap3 SNPs, resulting in 1,166,145 SNPs in our final dataset.

All analyses of real data used Plink 2 for GWAS, with the top 10 genetic principal components, age, and sex used as covariates. We used SumHer to estimate genetic covariances between feature and target phenotypes. Following the advice of Speed and Balding [12], we estimated genetic correlation with the LDAK-Thin model and heritability using the BLD-LDAK model, using tagging files provided at https://dougspeed.com/pre-computed-tagging-files. As a final QC step, we removed all estimates that were outside the range of possible true values (i.e. heritability outside [0, 1], genetic correlation outside [-1, 1], or standard error greater than two). This meant that each MaxGCP phenotype had a slightly different subset of the feature traits from which to select.

### 5.2 Details of the simulation

Realistic effect sizes and correlations among phenotypes are essential for evaluating MaxGCP. The PhenotypeSimulator R package [16] is designed for this purpose, and it can simulate phenotypes using real genetic data. We used this package to generate phenotypes with known effect sizes and correlations. To construct the cohort for this simulation, we randomly sampled 10,000 (White British, QC-passing) individuals and 100,000 (imputed, HapMap3, QC-passing) variants from the UK Biobank.

PhenotypeSimulator has many tunable parameters that control the genetic and phenotypic properties of the simulated phenotypes. Heritability, genetic correlation, and phenotypic correlation were the only relevant parameters for this study. We took a two-step approach to setting these parameters. First, we picked two sets of values that we believed to be realistic. Second, to evaluate the sensitivity of our results to these choices, we individually varied each parameter while keeping the others constant at realistic values.

To determine realistic settings for these parameters, we estimated the heritability, genetic correlation, and phenotypic correlation of real diseases in the UK Biobank. For convenience, we used only the 20 most common ICD-10 codes in our QC-passing White British cohort. After estimating these values in real data and performing QC on the results, we found that the median heritability was 0.05, the median genetic correlation was 0.49, and the median phenotypic correlation was 0.10. We sought to simulate phenotypes that approximately matched these values. PhenotypeSimulator input parameters define distributions, not the actual values of the resulting phenotypes. Accordingly, we adjusted the PhenotypeSimulator parameters so that the correlations match (Figure S1). We set heritability to 0.01, genetic correlation to 0.5, and phenotypic correlation to 0.0.

This simulation resulted in individual-level genotypes, phenotypes, and phenotypic genetic components. The format and structure of these data are very similar to real data, with a few notable exceptions. First, the simulated variant effect sizes are independent of one another, and there is no relationship between variant effect size and LD. This means that LDSC-like methods for heritability estimation are inapplicable. Second, we did not generate any covariates or confounders, as they are not relevant to the current analysis. Our simulation is intentionally much smaller than the UK Biobank dataset, with fewer variants, samples, and phenotypes. Our goal was not to recreate the UK Biobank in simulation, but to evaluate the feasibility of MaxGCP, the correctness of our definitions, derivation, and implementation, and whether MaxGCP achieves its aims.

**Figure S1:**
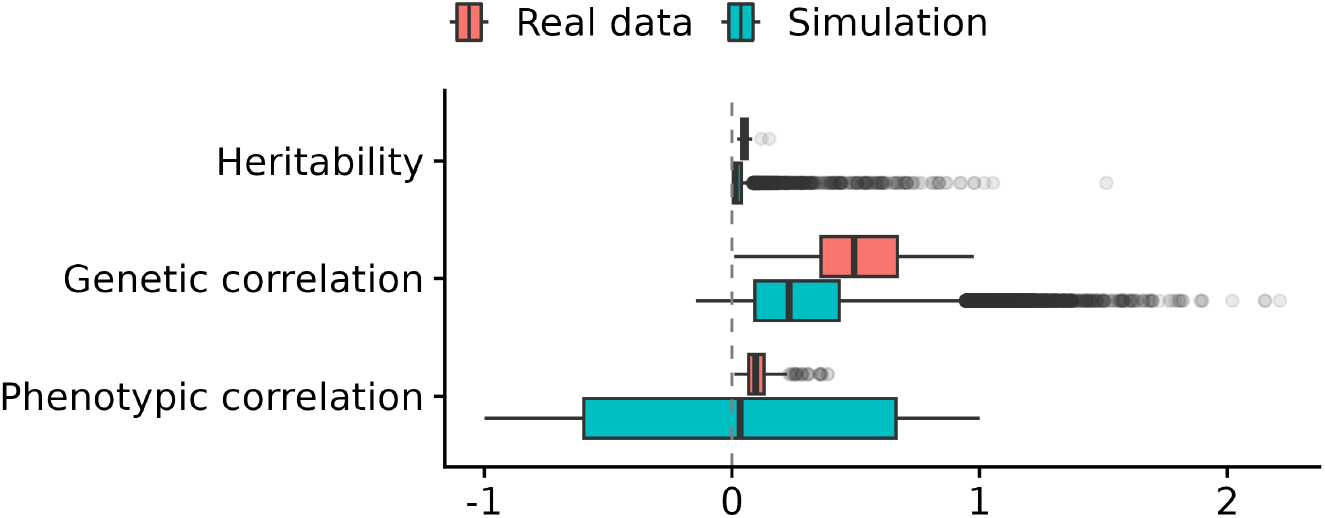
Distributions of heritability and correlation in real and simulated data. Shown is a comparison between simulation and the 20 most common ICD-10 codes in the UK Biobank, in terms of heritability, genetic correlation, and phenotypic correlation. The simulated phenotypes were generated using PhenotypeSimulator with expected heritability 0.01, genetic correlation 0.5, and phenotypic correlation 0. These simulation parameters were chosen so that the resulting simulation is similar or more conservative than the real data shown.

We simulated phenotypes with these settings and computed the actual heritability, genetic correlation, and phenotypic correlations. The distributions of these values showed good correspondence to their distributions in real data (Figure S1). MaxGCP depends on correlations and genetic signals to function, so it is expected to perform better with larger values of these parameters. We set the values of the simulation parameters conservatively so that the resulting distributions have similar or smaller means than we observe in real data. Overall, these results suggest that the simulation is reasonably similar to real data, albeit with known genetic effects.

### 5.3 MaxGCP top coefficients in real data

**Table 3:**
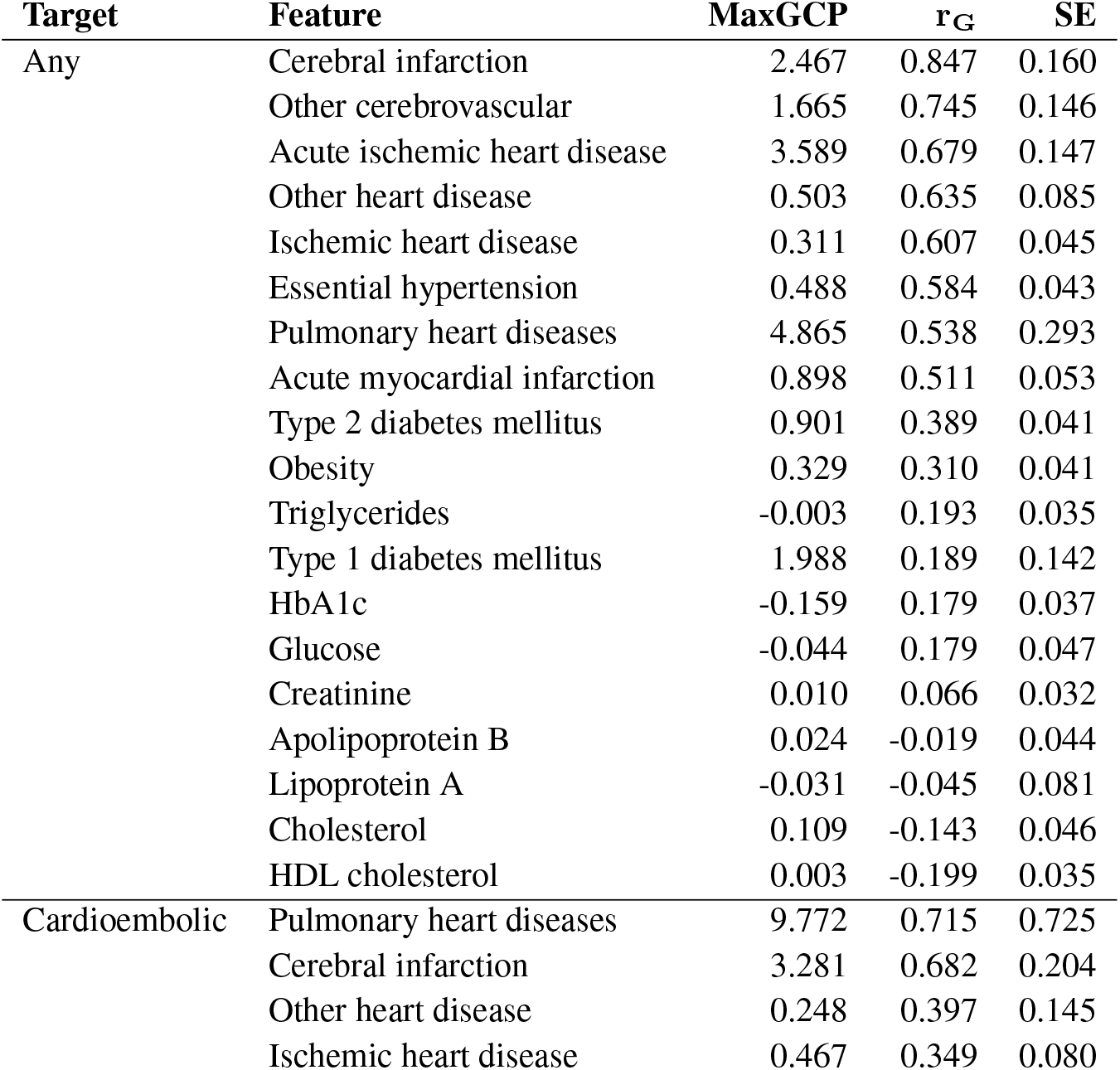
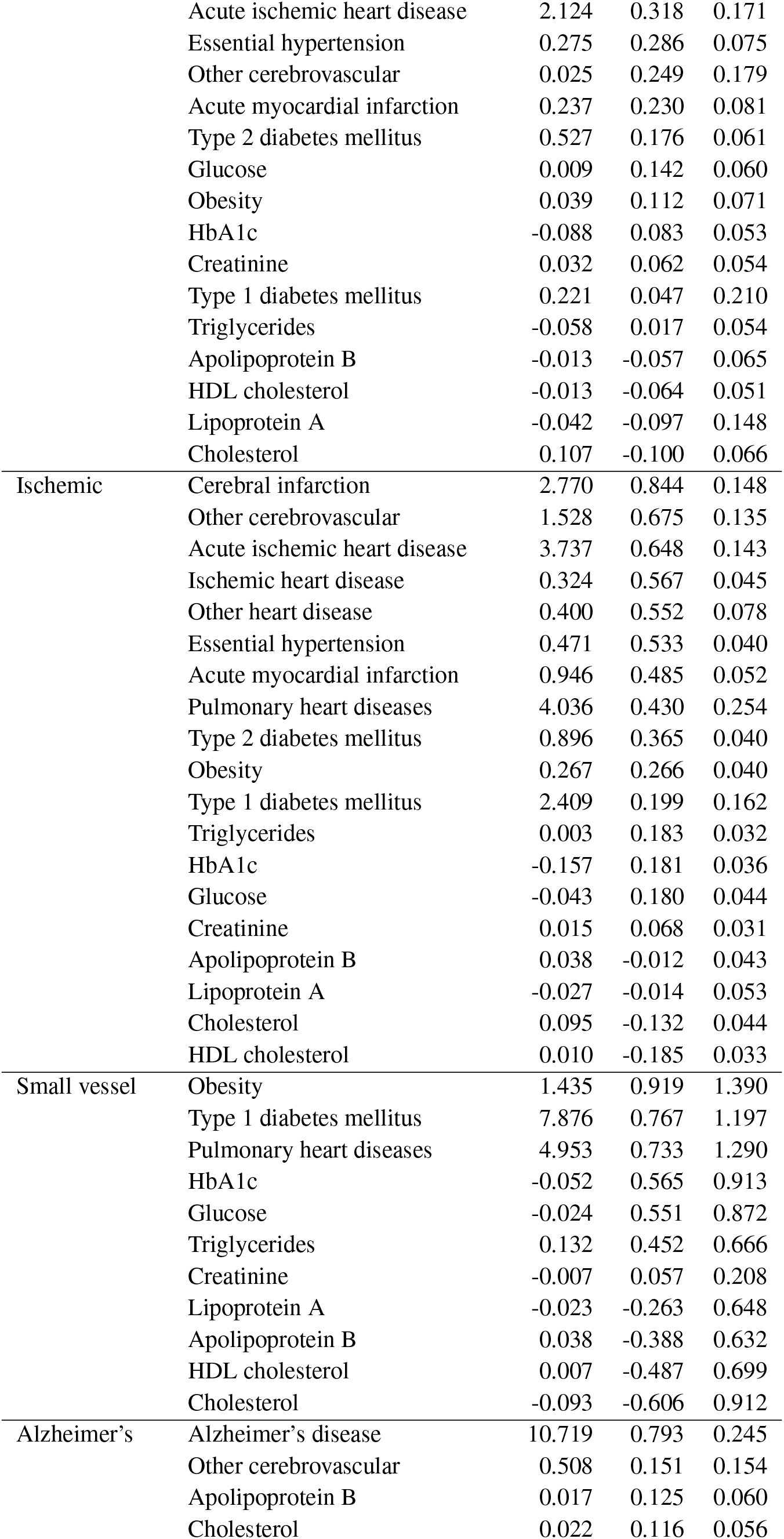
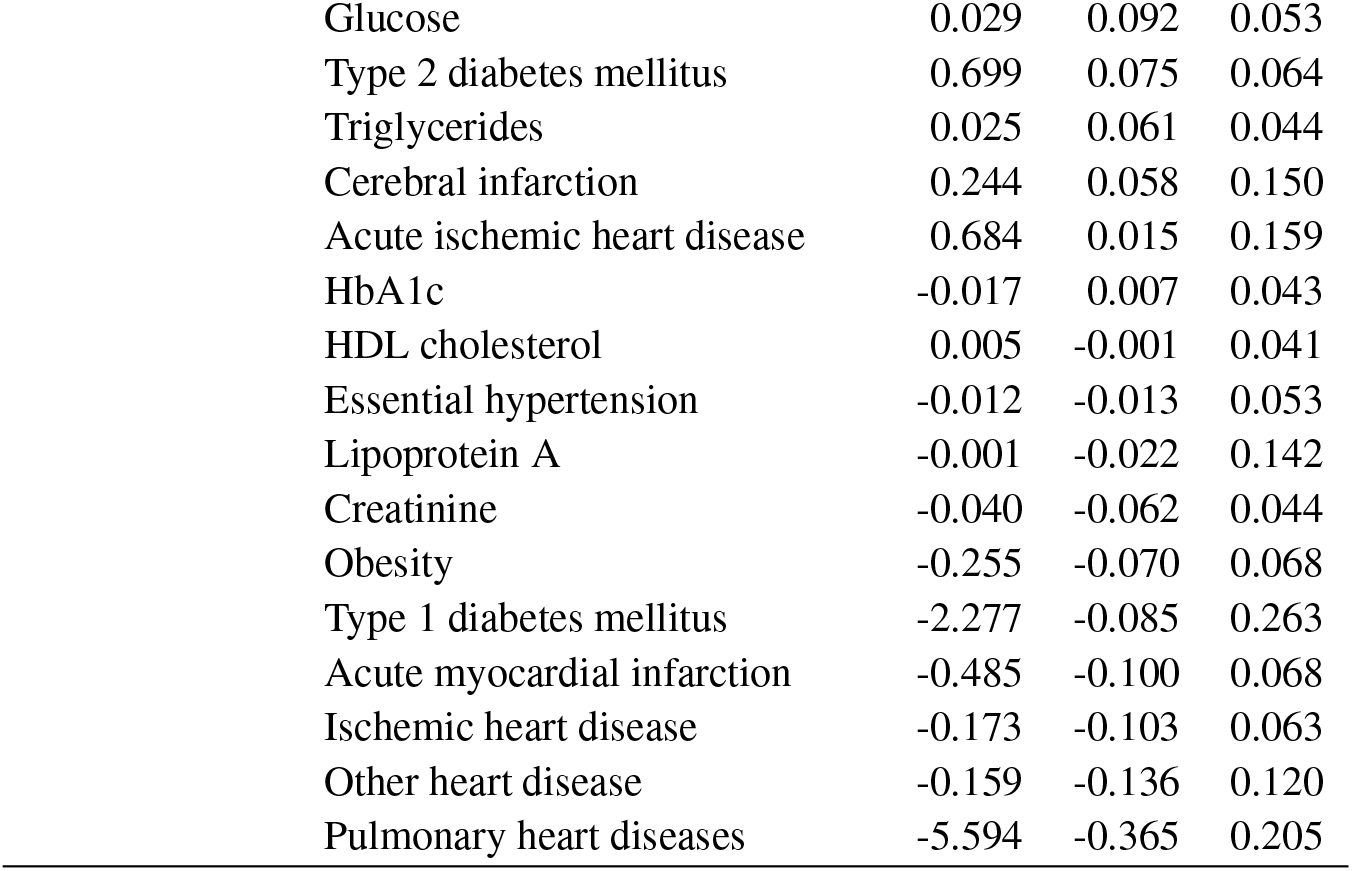
MaxGCP coefficients for target phenotype. Shown are selected features for each target phenotype. Each MaxGCP phenotype had a slightly different set of input features due to the QC process, in which noisy genetic covariance estimates were removed. Genetic correlations and standard errors were estimated using SumHer and the LDAK-Thin model.

### 5.4 Manhattan plots of GWAS results

**Figure S2:**
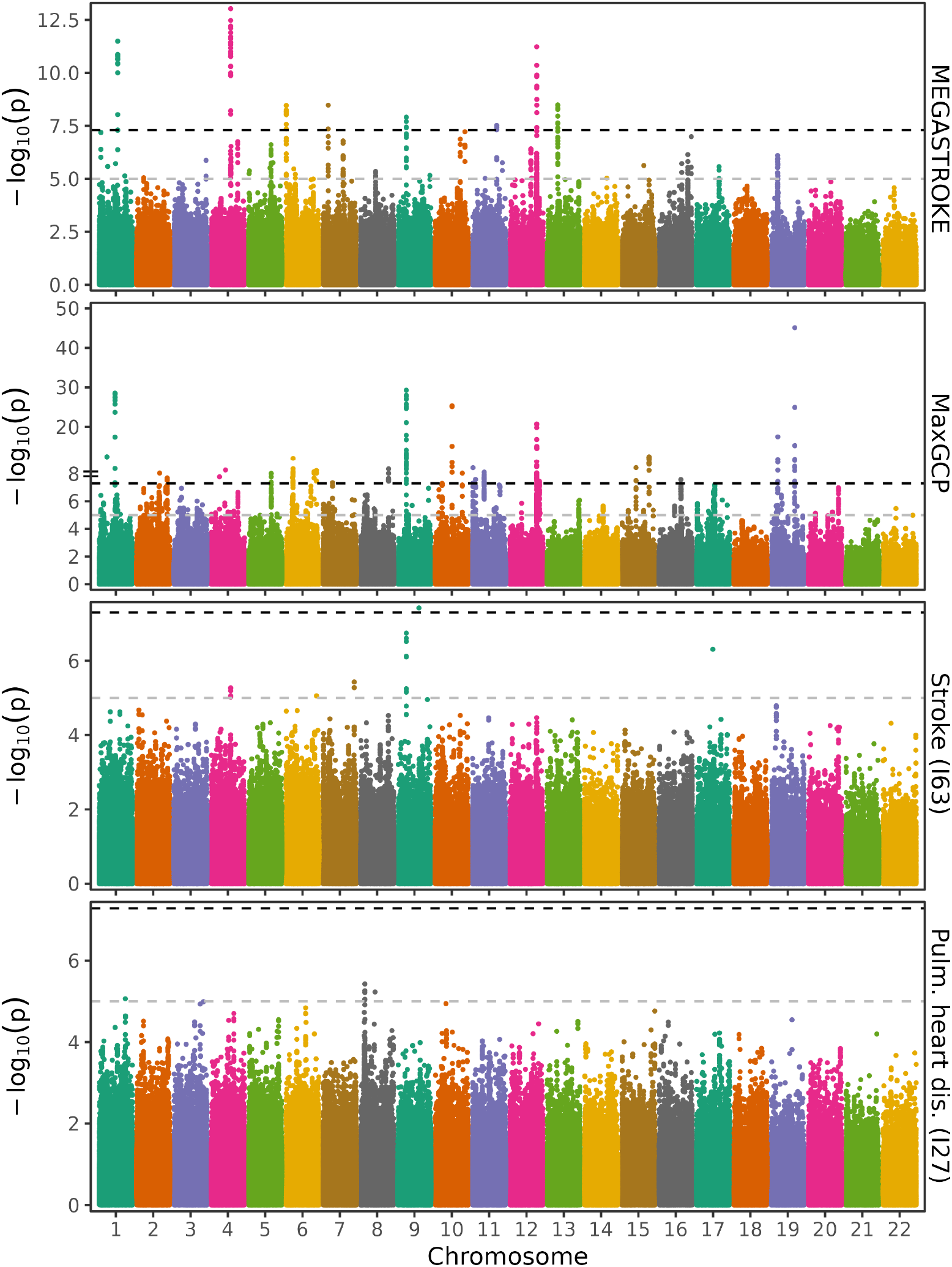
Comparison of GWAS results for stroke. Shown are GWAS summary statistics for the target, MaxGCP phenotype, naive phenotype, and top feature for the MEGASTROKE “Any stroke” phenotype. MEGASTROKE shows the MEGASTROKE “Any stroke” GWAS summary statistics. MaxGCP shows the corresponding MaxGCP phenotype. Stroke shows the the naive equivalent phenotype, defined as a single occurrence of any ICD-10 code starting with I63. The final facet shows GWAS summary statistics for pulmonary heart disease, the top feature in MaxGCP besides stroke itself.

**Figure S3:**
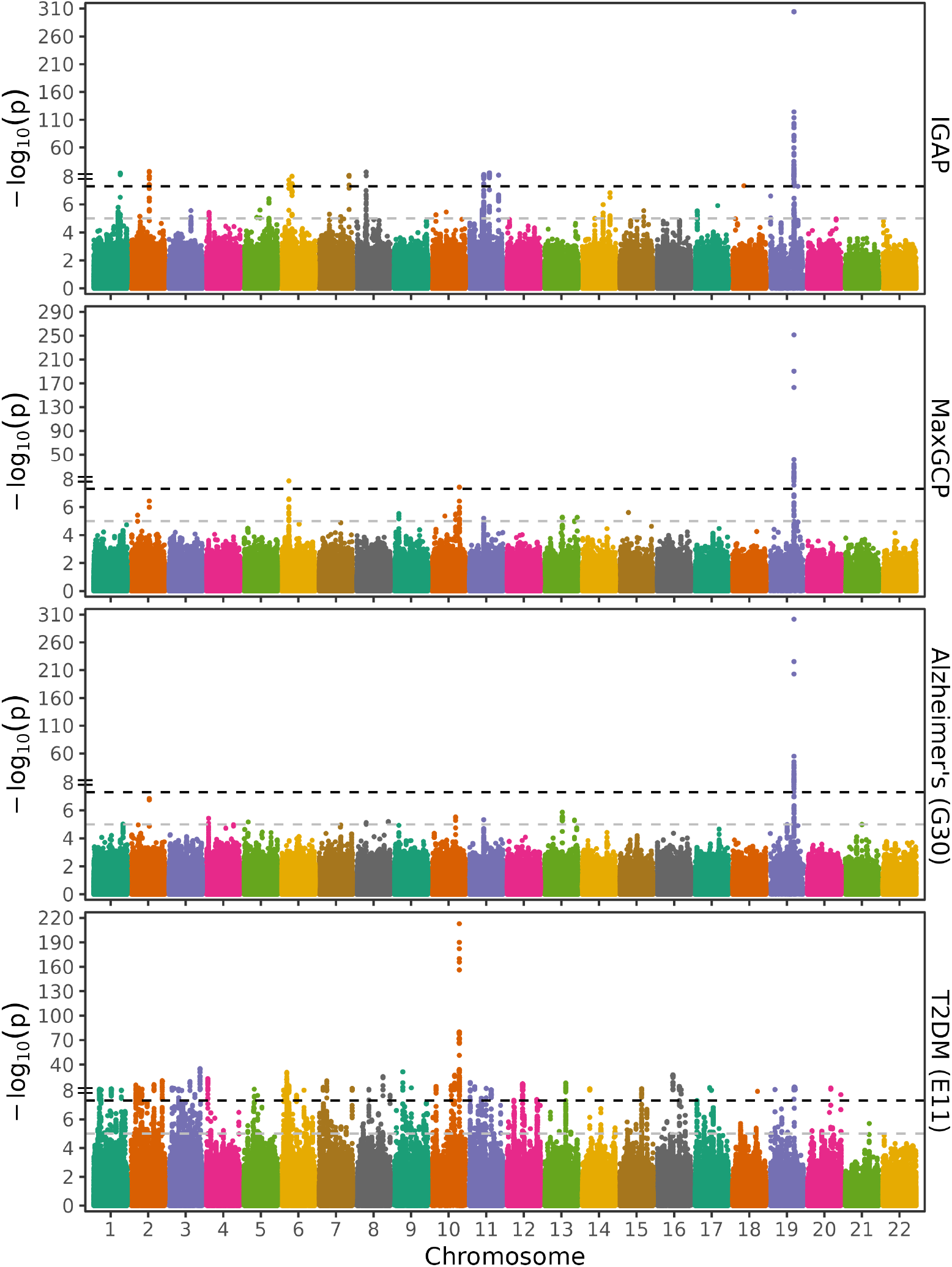
Comparison of GWAS results for Alzheimer’s disease. Shown are GWAS summary statistics for the target, MaxGCP phenotype, naive phenotype, and top feature for the IGAP Alzheimer’s disease phenotype. IGAP shows the IGAP GWAS summary statistics directly. MaxGCP shows the corresponding MaxGCP phenotype. Stroke shows the the naive equivalent phenotype, defined as a single occurrence of any ICD-10 code starting with G30. The final facet shows GWAS summary statistics for type 2 diabetes mellitus, the top feature in MaxGCP besides Alzheimer’s disease itself (ICD-10 code G30).

